# The proximal proteome of 17 SARS-CoV-2 proteins links to disrupted antiviral signaling and host translation

**DOI:** 10.1101/2021.02.23.432450

**Authors:** Jordan M Meyers, Muthukumar Ramanathan, Ronald L Shanderson, Laura Donohue, Ian Ferguson, Margaret G Guo, Deepti S Rao, Weili Miao, David Reynolds, Xue Yang, Yang Zhao, Yen-Yu Yang, Yinsheng Wang, Paul A Khavari

**Affiliations:** Program in Epithelial Biology, Stanford University, Stanford, CA 94305, USA; Program in Cancer Biology, Stanford University, Stanford, CA 94305, USA; Program in Genetics, Stanford University, Stanford, CA 943054, USA; Program in Biomedical Informatics, Stanford University, Stanford, CA 94305, USA; Department of Chemistry, University of California, Riverside, CA, 92521, USA

**Keywords:** SARS-CoV-2, BioID, proteomics

## Abstract

Viral proteins localize within subcellular compartments to subvert host machinery and promote pathogenesis. To study SARS-CoV-2 biology, we generated an atlas of 2422 human proteins vicinal to 17 SARS-CoV-2 viral proteins using proximity proteomics. This identified viral proteins at specific intracellular locations, such as association of accessary proteins with intracellular membranes, and projected SARS-CoV-2 impacts on innate immune signaling, ER-Golgi transport, and protein translation. It identified viral protein adjacency to specific host proteins whose regulatory variants are linked to COVID-19 severity, including the TRIM4 interferon signaling regulator which was found proximal to the SARS-CoV-2 M protein. Viral NSP1 protein adjacency to the EIF3 complex was associated with inhibited host protein translation whereas ORF6 localization with MAVS was associated with inhibited RIG-I 2CARD-mediated *IFNB1* promoter activation. Quantitative proteomics identified candidate host targets for the NSP5 protease, with specific functional cleavage sequences in host proteins CWC22 and FANCD2. This data resource identifies host factors proximal to viral proteins in living human cells and nominates pathogenic mechanisms employed by SARS-CoV-2.

**Author Summary:** SARS-CoV-2 is the latest pathogenic coronavirus to emerge as a public health threat. We create a database of proximal host proteins to 17 SARS-CoV-2 viral proteins. We validate that NSP1 is proximal to the EIF3 translation initiation complex and is a potent inhibitor of translation. We also identify ORF6 antagonism of RNA-mediate innate immune signaling. We produce a database of potential host targets of the viral protease NSP5, and create a fluorescence-based assay to screen cleavage of peptide sequences. We believe that this data will be useful for identifying roles for many of the uncharacterized SARS-CoV-2 proteins and provide insights into the pathogenicity of new or emerging coronaviruses.

## INTRODUCTION

Coronaviruses comprise a diverse family of large positive-sense single stranded (+ss)RNA enveloped viruses that cause respiratory and gastrointestinal disease. In addition to common seasonal coronaviruses, a number of strains can cause severe disease, as seen in the Severe Acute Respiratory Syndrome (SARS-CoV-1) virus outbreak in 2003 (1), the Middle Eastern Respiratory Syndrome (MERS) virus outbreak in 2012 (2) and the 2019 outbreak of SARS-CoV- 2 (3). This viral family has large (26 to 32kb) genomes that encode tens of viral proteins. All coronaviruses have a similar organization consisting of a large open reading frame encoding two overlapping polyproteins, ORF1A and ORF1B. These polyproteins are cleaved by one of two viral proteases, NSP3 and NSP5, with the resulting protein products sequentially numbered NSP1-NSPX. ORF1AB is invariably followed by structural genes, including the Spike protein (S), Envelope protein (E), Membrane protein (M), and Nucleocapsid protein (N). SARS-CoV-2 encodes NSP1-16 as well as the accessory proteins ORF3a, ORF3b, ORF6, ORF7a, ORF7b, ORF8, ORF9b, ORF9c, ORF10, and ORF14, though it is not known if each open reading frame encodes for a functional protein product. The function of many SARS-CoV-2 accessory proteins is either unknown or highly variable across differing coronaviruses, underscoring the need to begin mapping their putative localizations and functions.

Proximity proteomics (BioID) uses enzymes, such as the modified bacterial biotin ligase, BirA, to biotinylate nearby proteins on lysine residue-containing proteins within a radius of 10-20nm (4). When fused to a protein of interest it labels not only proteins that directly bind the fused protein but also those adjacent to it, enabling rapid isolation of biotinylated proteins whose identity can provide clues about the localization and function of the protein studied. When coupled to mass spectrometry it provides an alternative to traditional tandem affinity purification and mass spectrometry (TAP-MS) (5). Whereas, TAP-MS can isolate protein complexes that stably bind the protein of interest in a manner robust enough to survive protein extraction, BioID-MS labels both transient and stable interactors in living cells, particularly those stabilized by cellular membranes that can be destroyed in traditional TAP-MS experiments. In this way, BioID may localize the cellular “neighborhoods” of a given fusion protein. We recently generated a biotin ligase derived from *Bacillus subtilis*, which has 50 times greater activity than the original *E. coli* BirA (4, 6), allowing decreased labeling times and increased signal-to-noise ratios. Applying proximity proteomics to SARS-CoV-2 viral proteins in human cells may facilitate insight into their localization and putative functions.

The actions of specific SARS-CoV-2-encoded proteins are only partially understood at present. The replication transcription complex, which includes the RNA-dependent RNA polymerase and other factors, and the structural proteins, which are necessary for protecting the newly synthesized genomes and assembling the viral particles, comprise the core viral replication machinery. Other viral gene products, generally termed accessory factors, are believed to be dedicated to manipulating the host environment to foster viral replication (7). One of the main functions of accessory factors is to block host antiviral response (8). Non-SARS-CoV-2 coronaviruses have also been shown to block host translation (9, 10), inhibit interferon signaling (11, 12), antagonize viral RNA sensing (13, 14), and degrade host mRNAs (15). The degree of homology between SARS-CoV-2 and other coronaviruses, suggests the existence of both shared and divergent host protein interactions between its viral proteins and those of the other members of the coronavirus family.

Here we used proximity proteomics to identify the human proteins vicinal to 17 major SARS-CoV- 2 proteins and, from that data and validation studies, to predict their likely location and function. We examined the intersection of the resulting atlas of human factors adjacent to SARS-CoV-2 viral proteins with risk loci associated with severe COVID-19 by genome wide association studies (GWAS). This nominated specific, viral protein-adjacent host candidates whose natural variation in expression may contribute to differences in COVID-19 susceptibility in the population. We also demonstrated that multiple SARS-CoV-2 products can affect host translation and host innate immune signaling and define a list of potential host targets and pathways for the NPS5 protease. Taken together, these resource data plot the location of the 17 major SARS-CoV-2 within the cell, define an atlas of human host proteins adjacent to them, and offer insight into potential pathogenic mechanisms engaged by SARS-CoV-2.

## RESULTS

### Host proteins proximal to viral proteins and their subcellular localization

To identify the human host proteins vicinal to the 17 major SARS-CoV-2 encoded viral proteins, HA epitope tagged fusions of BASU-BirA (6) were generated with each of these 17 viral ORFs (**Fig. 1A**). BASU was introduced at the N and C terminus to minimize disruption as previously described (16). Samples were prepared from plasmid-transfected 293T cells after 2 hours of biotin labeling and the biotinylated proteins were then isolated using streptavidin. Samples were divided for LC-MS/MS and immunoblotting (**Fig. S1**). MS data search was performed and protein lists were analyzed and scored using the Significance Analysis of Interactome (SAINT) method (17). Using a cutoff of a SAINT score of 0.9 generated a list of 2422 host proteins (**Fig. 1B, Fig. S2, Table S1**) across the 17 viral proteins studied, 514 of which were unique to a specific viral protein. These data (**Table S2**) comprise a compendium of candidate human proteins adjacent to SARS-CoV-2-encoded proteins.

**Fig. 1.**
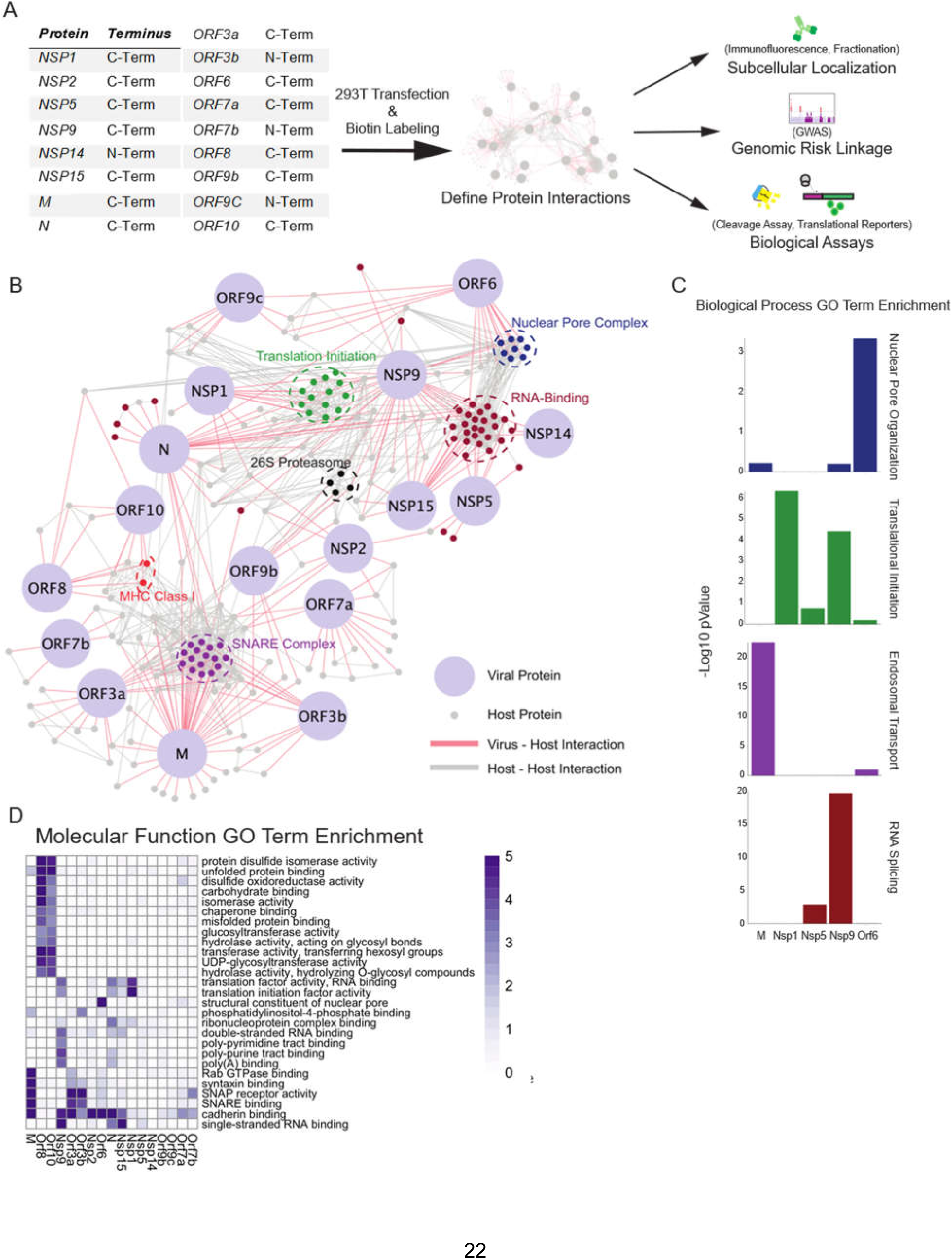
Proximal Interactome of 17 SARS-CoV-2 proteins. **A)** Schematic of BioID workflow. **B)** Curated network of SARS-CoV2 virus-host protein associations (SAINT ≥ 0.9) obtained from BASU BioID. Coronavirus proteins are labeled in light blue and virus-host interactions are connected by red edges, while host-host protein interactions obtained from high confidence STRING interactions are labeled in grey. Highlighted node clusters of similar function, including 26S proteasome components (black), MHC Class I (red), nuclear pore (dark blue), RNA-binding (maroon), SNARE complex (purple) , translation initiation complex (green) proteins were selected based on GO term analysis. **C)** Selected biological process GO term enrichment; enrichment scores are given as -Log10 p-values. Selected GO terms are nuclear pore organization, translational initiation, endosomal transport, and RNA splicing. **D)** Heatmap of molecular function GO term enrichment of SARS-CoV-2 proteins. All presented GO terms have a -Log10 p-value >3 for the Nucleoprotein, the listed non-structural proteins, or the listed open reading frames or a -Log10 p-Value >5 for the M membrane protein.

The identity of these 2422 human proteins provided clues to SARS-CoV-2 biology. Molecular function analysis (**Fig. 1B-D**) identified processes associated with SARS-CoV-2 viral protein impacts. This included translation initiation, RNA binding, the 26S proteasome, signaling, and SNARE-associated intracellular transport. It also identified adjacencies to major histocompatibility (MHC) proteins and components of the nuclear pore complex (NPC). A number of these processes, such as protein translation, are known processes affected by coronaviruses, while others, such as RNA-binding, are less well characterized.

To begin to map putative localizations for the 17 studied SARS-CoV-2 proteins within the cell, cellular component GO-term enrichment analysis was performed (**Fig. 2A**), which pointed to possible intracellular localizations for each viral protein based on curated knowledge of the host proteins identified adjacent to each viral protein. To validate and extend this, protein fractions were prepared from cells expressing each SARS-CoV-2 protein studied. These included four overlapping fractions: a) cytoplasm b) cytoplasm/membrane c) nucleus/membrane, and d) nucleus (**Fig. 2B**). Integrating GO-term analysis with immunoblotting of these fractions enabled predictions of the likely intracellular localization of each viral protein (**Fig. 2C**). We further confirmed NSP5 diffuse expression and ORF3a membrane localization through immunostaining. Many SARS-CoV-2 accessory proteins concentrate in the ER or in ER-proximal membranes (M, ORF3a, ORF3b, ORF6, ORF7a, ORF7b, ORF8, and ORF10). A number, however, appear to be predominantly cytoplasmic (NSP1, NSP2, NSP5, NSP9, NSP15, ORF9b) and, interestingly, several appear to localize in part to the nucleus (NSP14, ORF6, ORF9c). The localization predicted from these data is consistent with observations from other recent work (16, 18). Of the membrane localized proteins, subtle differences in location could be inferred. In the case of M protein, association with membranes in the endocytic pathway as well as lysosomal membranes was predicted. ORF8 and ORF10 clustered similarly with enrichment for ER interactions in the lumen. These data indicate that specific SARS-CoV-2 may display increased localization to a variety of intracellular sites, including the cytoplasm, nucleus and distinct endomembranes.

**Fig. 2.**
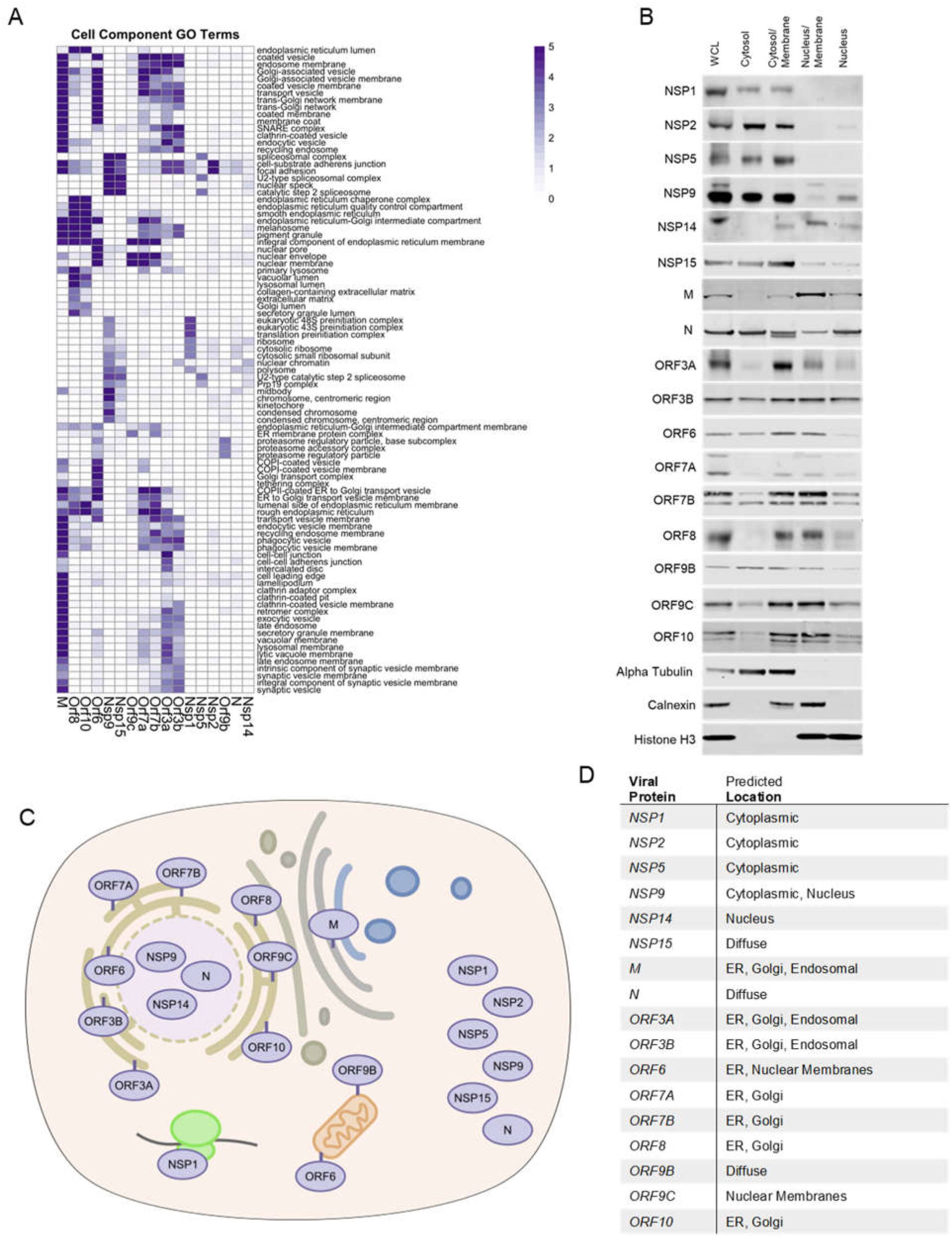
Localization of SARS-CoV-2 proteins. **A)** Heatmap of cell component GO term enrichment of SARS-CoV-2 proteins. **B)** Western blots of SARS-CoV-2 viral protein-expressing HEK293T cell fractions; whole cell lysate (WCL), cytosol, cytosol/membrane, nucleus/membrane and nucleus fractions. Alpha-tubulin, calnexin, and histone H3 were used as fractionation controls for cytosol, membrane, and nucleus respectively. Schematic **C)** and table **D)** depicting the predicted location of all SARS-CoV-2 proteins surveyed in this study based on both the BioID and fractionation analysis.

### Viral proximal interactors include drug targetable host genes

There is a lack of SARS-CoV-2 specific antiviral therapies or against coronaviruses generally. Many current and experimental therapeutics were developed for activity against other viruses and are being tested for cross efficacy against SARS-CoV-2. Others are therapies known to have broad antiviral effects. There is significant interest in developing drugs that directly target SARS-CoV-2 viral proteins, but research and development may take years before use in patients. Another approach is using drugs against host genes critical to virus infection and replication. For example, drugs targeting ACE-2, the main receptor for SARS-CoV-2, or ACE-2 expression and function have been pursued. To expand the list of possible drugs beyond entry inhibitors, we compared the viral proximal proteome generated in this study against the “druggable” genome, which include databases of the gene targets of available drugs. This generated a list of 47 host genes (**Fig. S3, Table S3**) and highlights, as previously reported (16), a group of cellular kinases associated with N protein. The viral nucleocapsid has been shown to be phosphorylated and phosphorylation is suggested to be important for its function (19, 20). This highlights cellular kinase inhibitors as drugs with possible activity against SARS-CoV-2.

### GWAS-linked host proteins in the viral proximal proteome

The genetic basis for the wide spectrum of COVID-19 severity in different individuals within the human population is not fully understood. A number of recent genome wide association studies (GWAS) studies have endeavored to map genetic risk loci associated with SARS-CoV-2 infection and COVID-19 clinical severity (21, 22). These studies leverage large numbers of patients to identify SNPs that are correlated with outcomes such as infection and severity of disease, including hospitalization and mortality. Such linkage studies have identified a number of non-coding variants that may perform a regulatory function, for example, by altering expression of effect genes (eGenes) important in host susceptibility to SARS-CoV-2.

To determine if any putative COVID-19 risk-linked regulatory variants might control the expression of host proteins proximal to SARS-CoV-2 viral proteins, the following analysis was performed. Using publicly available data from GWAS studies (21, 22), all single nucleotide polymorphisms (SNPs) associated with increased risk of COVID disease that reside in noncoding DNA were identified. These were filtered for variants localized to open chromatin, characteristic of regulatory DNA, in cell types relevant to COVID-19 pathogenesis, including immune and pulmonary cells. The resulting disease risk-linked variants were further distilled to those identified as expression quantitative trail loci (eQTLs) for specific putative eGene targets (**Fig. 3A**). These eGenes, which represent a set of genes whose expression may be controlled by natural variants in the human population linked to COVID-19 risk, were then intersected with the atlas of host factors identified as adjacent to SARS-CoV-2 viral proteins by proximity proteomics. Publicly available protein interaction data was then integrated to project the connectedness of resulting gene set (**Fig. 3B**). The resulting network was notable for host proteins implicated in cytokine signaling, cell cycle control, transcription, and translation, suggesting that genetic susceptibility to COVID-19 may link to variations in the expression of proteins that mediate these processes.

**Fig. 3.**
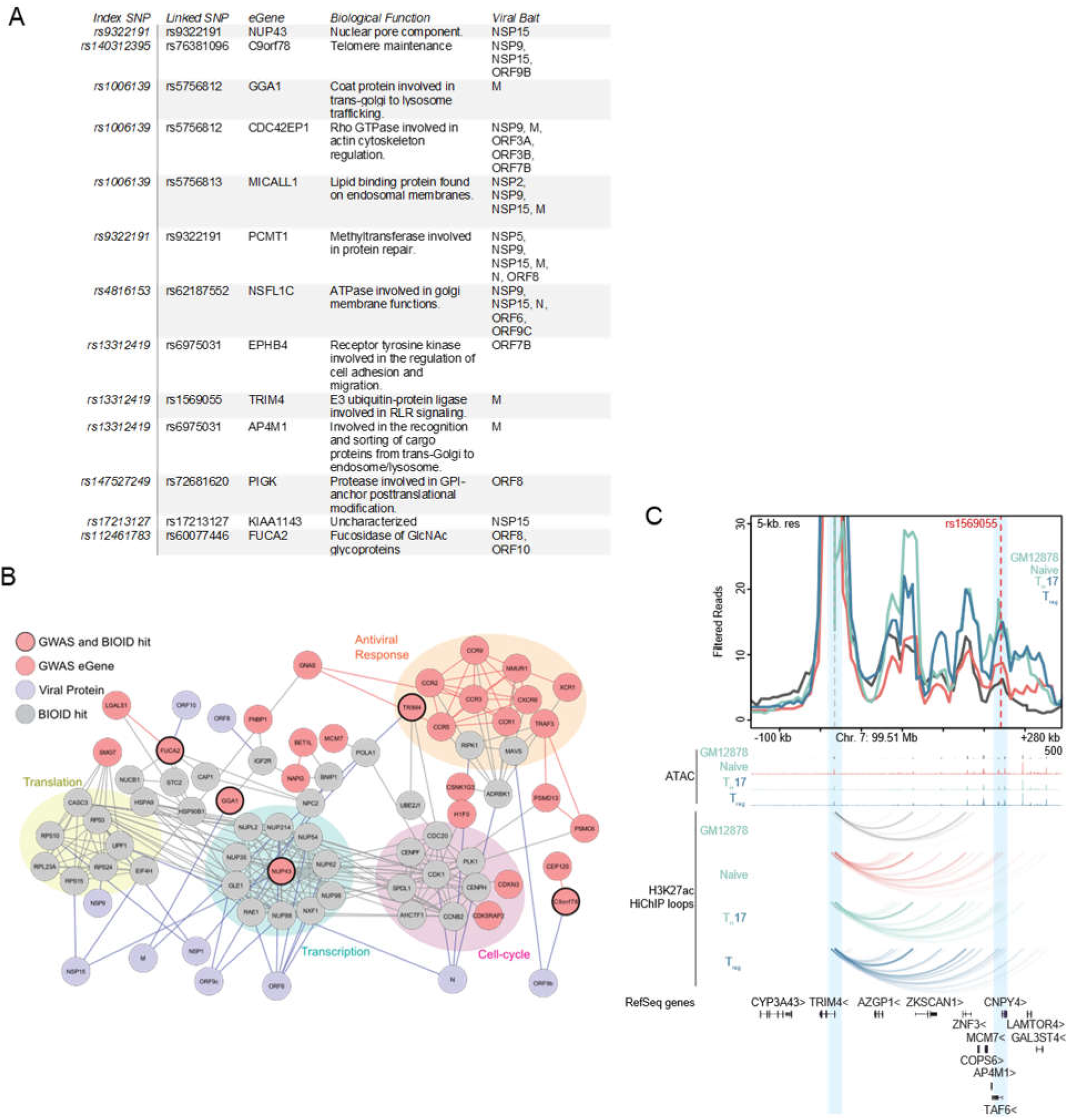
COVID disease risk eGenes proximal to viral proteins. **A)** Table of GWAS risk SNPs which also scored as BioID hit. **B)** Map of connectedness of eGenes (Mauve) with BioID interactors (Gray) and the corresponding viral proteins (Purple). eGenes also identified by BioID are outlined in black. GWAS-identified eGenes-associated with antiviral response, cell-cycle, transcription, and translation are also highlighted. **C)** Virtual 4-C plot showing chromatin contact between TRIM4 promoter and linked COVID disease risk SNP (rs1569055). Genome tracks showing ATAC peaks and contact loops of GM12878, Naïve T, Th- 17, and T-reg cells.

Among proteins identified by this analysis was TRIM4, a RING E3 ligase, that activates type I interferon signaling through activation of the cytosolic RNA sensor RIG-I. TRIM4 was significantly associated with SARS-CoV-2 M protein in proximity proteomics data (**Table S1**) and, using eQTLgen (23), a regulatory SNP (rs1569055) approximately 230kb downstream of the TRIM4 promoter was recently associated with increased COVID severity in patients (22). High-C chromatin immunoprecipitation (HiChIP) data from the immortalized B-cell line GM12878 as well as primary T-cell populations demonstrated chromatin looping from the SNP to the TRIM4 promoter (**Fig. 3C**). Looping of this SNP increased contact strength in naïve, T-regs and Th-17 T-cells. TRIM4 is one of a group of ubiquitin ligases (24–27) that can activate RIG-I during RNA-sensing and subsequent antiviral signaling. Altered expression coupled with disruption by potential association with the SARS-CoV-2 M protein supports a model with the following features; a) individuals with this regulatory variant may express less TRIM4 b) physical association with SARS-CoV-2 M protein further reduces functional TRIM4 c) a relative reduction in biologically active TRIM4 leads to reduced innate immune signaling d) this reduction leads to increased susceptibility to SARS-CoV-2 pathogenesis. Integrating proximity proteomics data with genetic risk eQTL variants may help identify such candidate susceptibility mechanisms for natural variations in disease outcomes within the population.

### Predicted viral antagonism of host protein translation and antiviral response

NSP1 is a part of the viral polyprotein ORF1 and during normal viral replication is cleaved and liberated by the viral protease NSP3. Earlier work has identified NSP1 of SARS-CoV-1 as a potent inhibitor of translation in a mechanism that involves interactions with the host ribosomes (9, 12). Recently other groups have shown that NSP1 of SARS-CoV-2 similarly blocks translation through interaction with the 40s ribosome (28, 29). High confidence proteins proximal to NSP1 included EIF3A, EIF3B, EIF2G, and EIF4G2 (**Fig. 4A**) of which the first 3 are components of the EIF3 translation initiation complex. Interestingly, members of the EIF3 complex were not identified as high-confidence interactors by traditional TAP-MS studies(16, 18). To test if SARS-CoV-2 NSP1 inhibits host translation, NSP1 was expressed in HEK293T cells followed 24 hours later by transfection of in-vitro transcribed capped and polyadenylated mRNA expressing luciferase. NSP1 reduced luciferase signal by (49.7%) as compared to GFP control (**Fig. 4B**), demonstrating that NSP1 can inhibit host cap-dependent translation, consistent with data reported by others (28, 29). To determine if NSP1 could inhibit translation of host-derived 5’ UTRs and host IRES elements, two host UTRs (IFIT1 and ISG15) were subcloned separately upstream of luciferase along with two host IRES sequences (XIAP1 and APAF1) and luciferase measured in cells transfected with or without NSP1 construct. NSP1 reduced luciferase signal of both 5’ UTRs (IFIT1 = 55.2%, ISG15 = 53.1%) and IRES elements (XIAP1 = 55.0%, APAF1 = 40.0), indicating a block in translation of these elements (**Fig. 4B**). Lastly, NSP1 effects were tested on the SARS-CoV-2 5’ UTR and the Cricket Paralysis Virus (CRPV) IRES. CRPV IRES is a minimal viral-derived IRES that initiates translation completely independent of EIF3. Surprisingly, NSP1 blocked both viral elements (SARS-CoV-2 = 59.1%, CRPV = 52.2%) compared to GFP control. NSP1 therefore exhibits broad translation inhibition of mRNAs containing various regulatory elements, suggesting NSP1 action on the initiating ribosome, however, additional actions, such as mRNA cleavage (9, 15), may also be operative.

**Fig. 4.**
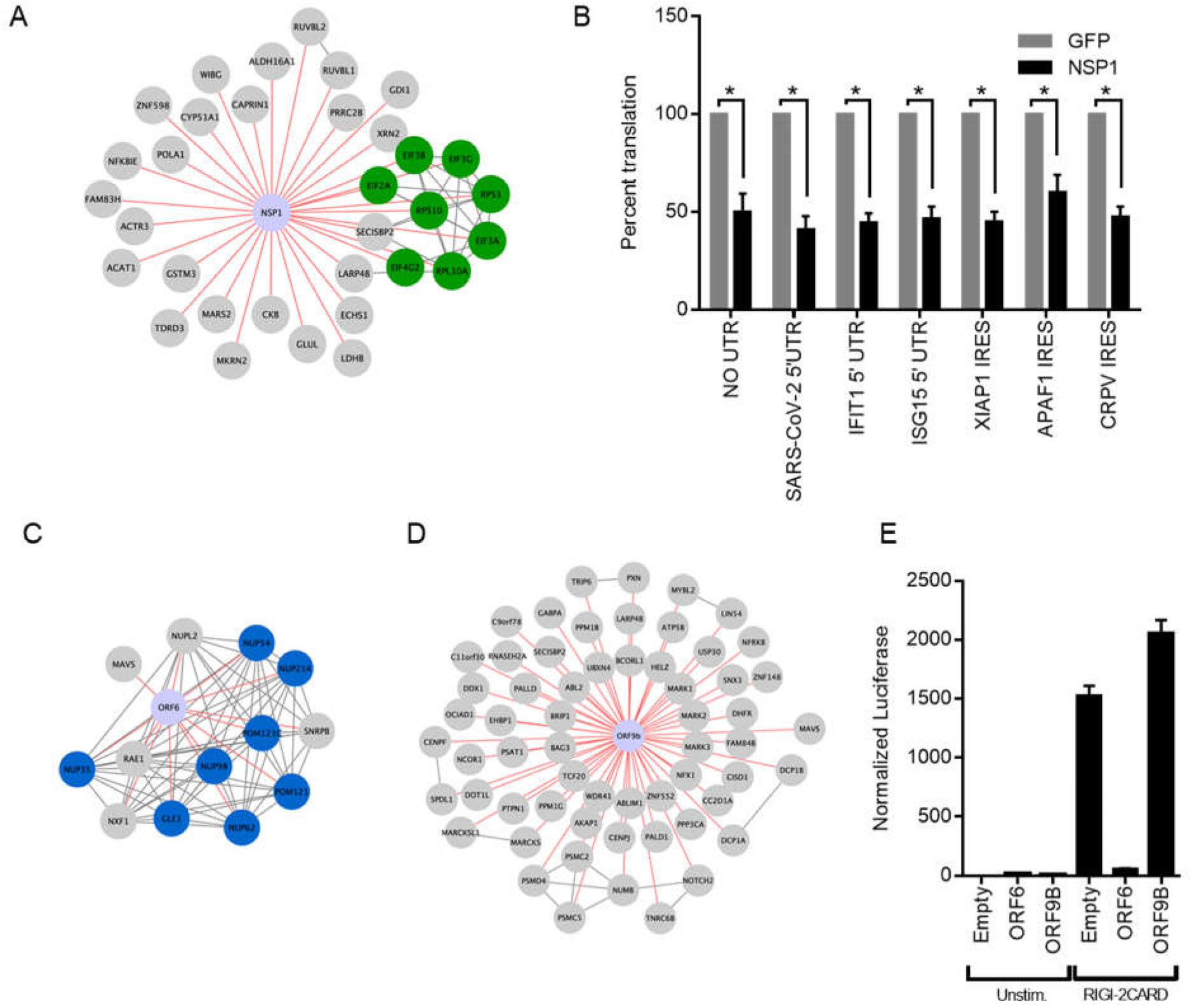
NSP1 and ORF6 disruption of host translation and innate immune signaling. **A)** Curated map of NSP1 proximal interactors. Highlighted are host proteins involved in translation initiation. **B)** Effect of NSP1 on translation of in vitro transcribed, capped polyadenylated transcripts containing 5’ UTRs from SARS CoV-2, *IFIT1*, and *ISG15* as well as IRES elements from *XIAP1, APAF1*, and CRPV. Data shown is the average of three independent experiments and significance was calculated using Student’s T Test where * indicates p value<0.005. Curated maps of ORF6 **C)** and ORF9b **D)** proximal proteins showing nuclear pore protein complex association with ORF6 and MAVS association with both ORF6 and ORF9b. **E)** Effect of ORF6 and ORF9b on *IFNB1* promoter activity after RIG-I 2-CARD induction. Normalized luciferase shown is the ratio of nano luciferase/firefly luciferase normalized to empty vector control. Data shown is the average of three independent experiments.

Host innate immune detection and signaling pathways are heavily targeted by viral proteins, especially accessory proteins (30). Mitochondrial Activation of Viral Signaling (MAVS) is a critical signaling adaptor for RIG-I like receptors (RLR) cytosolic sensing pathway (31–34). It recruits activated RLR sensors RIG-I and MDA-5 at mitochondrial and mitochondrial-proximal membranes and leads to the activation of both IRF3 and NF-κB and expression of type-I interferons (35). RIG-I and MDA-5 recognize various types of non-host or aberrant RNA species and are critical for host defense against RNA viruses (36). MAVS was found as a high confidence protein proximal to two SARS-CoV-2 proteins: ORF6 and ORF9b (**Fig. 4C-D**). ORF6 has been found to inhibit type-I interferon SARS-CoV-2 (37, 38) and the closely related SARS-CoV-1 (11, 39). One study demonstrated that ORF6 inhibition of type-1 interferon expression was linked to ORF6 binding to nuclear import complex RaeI/Nup98 (38), both of which were also captured as proximal interactors of ORF6, but not ORF9B. We tested the ability of our SARS-CoV-2 ORF6 and ORF9b constructs to inhibit RLR signaling by co-transfecting constructs expressing ORF6 or ORF9B along with a reporter expressing nanoluciferase under the control of the *IFNB1* promoter for interferon β1 along with a second reporter constitutively expressing firefly luciferase. To activate RLR signaling we transfected in a plasmid expressing a truncated version of RIG-I only containing the 2 CARD domains. This truncation is constitutively recruited to MAVS and initiates signaling in absence of any RNA stimulus and will test the viral proteins ability to block any signaling downstream of sensing. ORF6 significantly inhibited RIG-I 2CARD activation of *IFNB1* promoter activity by 96 percent (**Fig. 4E**) while ORF9b showed no effect on inhibiting *IFNB1* promoter activity. These data demonstrating ORF6 proximity to MAVS, along with ORF6 inhibition of *IFNB1* promoter induction, implicate ORF6 impairment of MAVS in the RLR innate immune signaling pathway.

### NSP5 proteomics prediction of potential host cleavage targets

NSP5 is one of two critical proteases encoded by SARS-CoV-2 and is also known as SARS-CoV- 2 3CLpro due to its similarity to picornavirus 3C proteases and a number of other +ssRNA viruses. These proteases all contain chymotrypsin-like folds and a triad of residues harboring the critical cysteine residue (40). 3CLpro-like proteases are considered important therapeutically since they are essential for cleaving large polyprotein products produced by +ssRNA viruses and chemical protease inhibitors may act broadly across members of a given virus family (41, 42). In addition to their necessity in the virus life cycle, many viral proteases can target host proteins and specifically affect antiviral responses or other cellular processes (43–45). Complementing previous efforts to infer targets of the NSP5 protease, we identified 34 host proteins in the NSP5 proximal proteome (**Fig S2C**). To nominate possible host targets of NSP5 whose levels are decreased upon protease expression, we performed SILAC mass spectrometry comparing wild type SARS-CoV-2 NSP5 to the catalytically-inactive NSP5^C145A^ mutant (16, 46). Residue 145 is the critical catalytic cysteine and mutation to alanine prevents protease activity (47). A number of host proteins showed significant depletion in cells expressing wild type NSP5, but not protease-inactive NSP5^C145A^ (**Fig. 5A**). Combining both data generated identified an additional 26 candidates resulting in a pool of 60 potential host protein targets for NSP5 (**Fig 5B**).

**Fig. 5.**
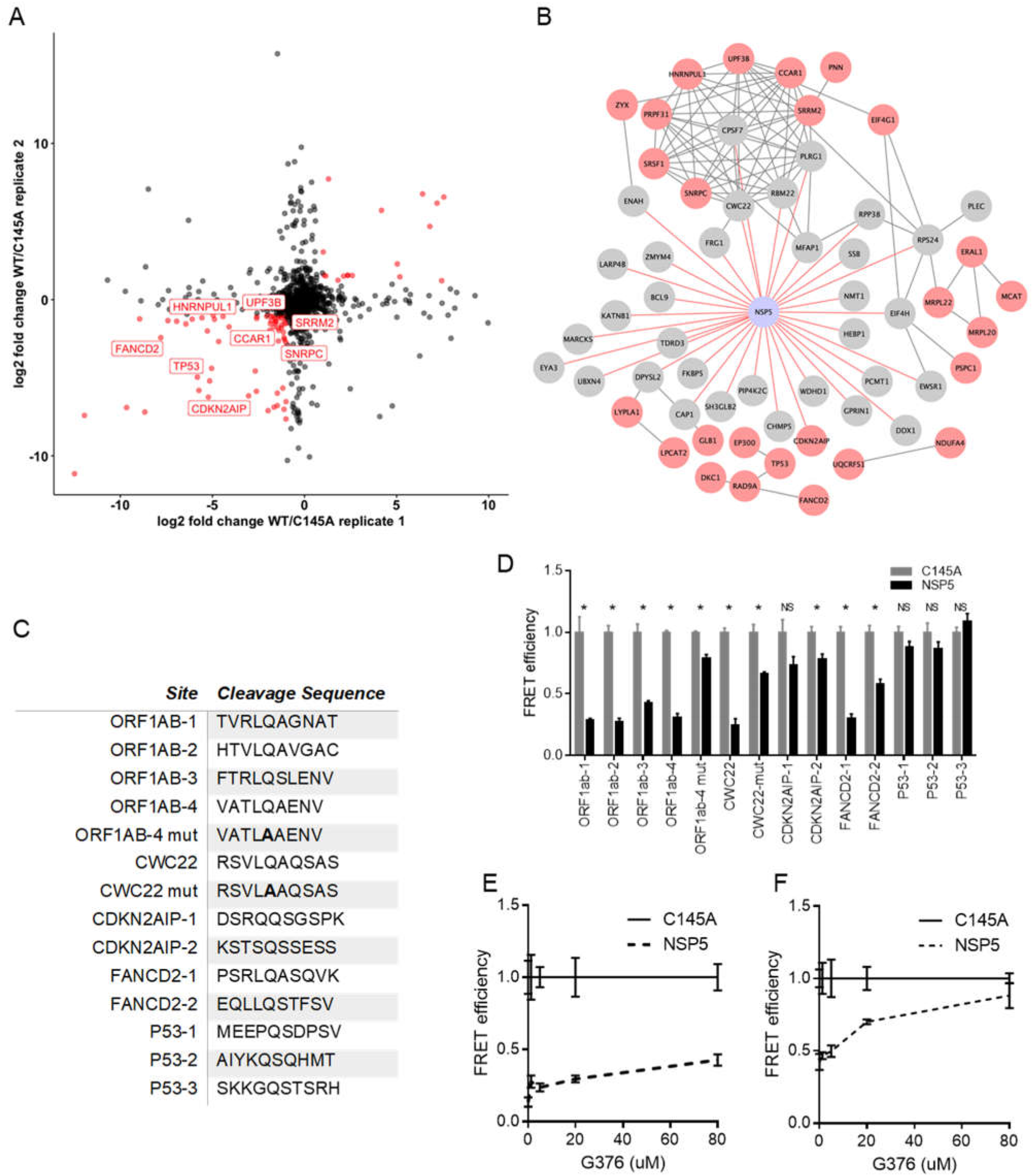
BioID and SILAC MS identify candidate targets for the viral protease NSP5. **A)** Comparison of two biological replicates of protein abundance in HEK293T cells expressing either NSP5 wild type (WT) or the catalytically inactive NSP5^C145A^ mutant by log_2_ fold change. **B)** Map of NSP5 proximal interactome (Gray) overlaid with host proteins decreased in abundance in SILAC (Red). CDNK2AIP was detected as both a BioID hit and decreased in abundance in SILAC. **C)** Peptide cleavage assay of four sequences from SARS-CoV-2 polyprotein ORF1AB (PP1ab) and the indicated host genes: CWC22, CDNK2AIP, FANCD2, P53. Normalized FRET signal is shown comparing HEK293T cells expressing either wild type NSP5 or NSP5^C145A^. ORF1ab and CWC22 mutant (mut) sequences contain QS→AS mutation in the peptide sequence. Data shown is representative of three independent experiments and significance was calculated using Student’s T Test. * indicates p value <0.05, NS not significant. Dose-dependent effect of coronavirus protease inhibitor G376 on cleavage of ORF1ab-2 **D)** and FANCD2-2 E) peptide sequences.

**Fig. 6.**
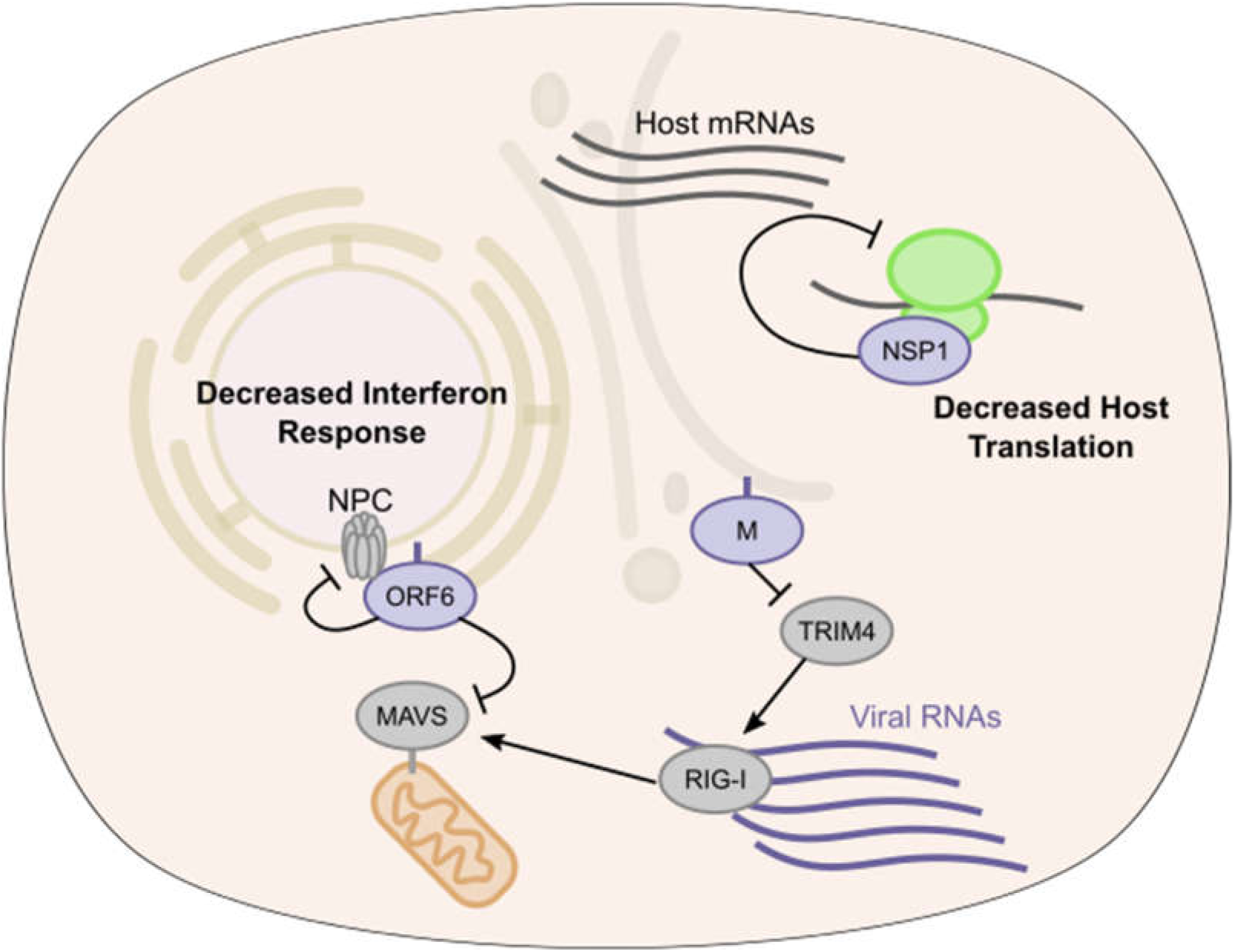
SARS-CoV-2 proximal proteins in translation and interferon activation. Model of SARS-CoV-2 antagonism of host antiviral response. ORF6 protein inhibits RLR signaling leading to decreased type I interferon and ISG transcription, M protein through TRIM4 interactions may also alter host response. NSP1 disrupts host translation of transcripts containing both ISG 5’ UTR and stress responsive IRES elements.

To begin to examine potential cleavage of these candidate proteins by NSP5, we searched their peptide sequences for potential cleavage sites using a published a cleavage prediction algorithm (48). We then took these peptide sequences and tested them for cleavage by NSP5 using a loss of fluorescence resonance energy transfer (FRET) fluorescence assay. In brief, potential cleavage sites were inserted between a FRET pair and then this construct was co-transfected along with plasmids expressing either NSP5 or the NSP5^C145A^, with loss of FRET signal only after wild type NSP5 expression as indicative of cleavage. Four sequences taken from SARS-CoV-2- ORF1AB polyprotein, which is normally cleaved by NSP5, were cleaved as expected and as demonstrated by loss of FRET signal (**Fig. 5C-D**). Testing of sequences from human CDKN2AIP, CWC22, FANCD2, and P53 proteins indicated NSP5 cleavage of one CWC22 and two FANCD2 peptide sequences (**Fig. 5C-D)**. Neither CDKN2AIP nor P53 sequences tested were cleavable by NSP5 in our assay and their depletion in the SILAC data may represent indirect effects of NSP5 activity. CWC22 is a component of the RNA spliceosome required for pre-mRNA splicing via promotion of exon-junction complex assembly (49, 50). FANCD2 is activated by ATM and localizes at BRCA1 foci during DNA damage (51). These data suggest that SARS-CoV-2 may target host RNA splicing and DNA damage pathways via NSP5-mediated reduction in key proteins, namely CWC22 and FANCD2, that are involved in these processes.

As noted, viral protease inhibitors are a powerful class of drugs that potently block viral replication by preventing processing of viral polyproteins into functional subunits. Inhibition of viral proteases should also prevent cleavage of host proteins which may serve to blunt toxic effects on infected cells. GC376 is a NSP5 protease inhibitor developed against feline coronavirus, the causative agent of fatal feline infectious peritonitis. Recent reports showed GC376 to be effective against SARS-CoV-2 NSP5. We tested the effect of GC376 on the cleavage of the ORF1ab-2 and FANCD2-2 peptide sequences. Using a range of concentrations up to 80 µM, ORF1ab-2 showed a modest inhibition of NSP5 as compared to NSP5^C145A^ in the FRET assay. FANCD2-2 showed a dramatic reduction in cleavage by NSP5 even at concentrations of 20 µM (**Fig. 5D-E**). These data support GC376 inhibition of SARS-CoV-2 NSP5 action on viral and human host protein sequences cleavable by the viral protease.

## DISCUSSION

Here we present a compendium of human host proteins adjacent to 17 SARS-CoV-2 viral proteins, with a goal to offer insight into potential mechanisms that these viral proteins may engage during pathogenesis. These data encompass the less well understood SARS-CoV-2 accessory factors and predict the localization of each these viral proteins as well as identify significant adjacencies to proteins that mediate core cellular processes, including translation, signaling, RNA interactions, and intracellular transport. For translation, SARS-CoV-2 NSP1 was found to be adjacent to subunits of the EIF3 translation initiation complex and proved a broad inhibitor of translation. For innate immune signaling, viral ORF6 was found proximal to the RLR pathway component, MAVS, with ORF6 potently inhibiting induction of the RLR downstream *IFNB1* promoter. Integration of GWAS data in COVID-19 identified SNPs associated with natural variation in the expression of specific genes, including the viral M protein-proximal TRIM4 activator of type I interferon, that may contribute to disease susceptibility differences in the human population. Comparing wild type NSP5 with its catalytically inactive point mutant helped identify proteins whose levels were decreased by this viral protease and nominated cleavage sequences in human CWC22 and FANCD2, implicating specific candidates for viral disruption of normal pre-mRNA splicing and DNA damage pathways, respectively. We also observed a number of SARS-CoV-2 proteins (M, NSP2, NSP9, NSP15, ORF6, ORF7a, ORF7b, ORF8, ORF9c, and ORF10) vicinal to nuclear pore proteins. Given that coronavirus replication takes place exclusively in the cytosol of cells, these interactions, if functional, might point to a viral role in disrupting nuclear import/export. This is further supported by several lines of genetic evidence. GWAS data suggest the importance of nuclear pore component NUP43 and overlap of our proteomics data with whole genome CRISPR screen hits from recently studies(52–54) suggests that the mRNA export factors MCM3AP and NXF1 are necessary for viral replication. Taken together, these data indicate potential intracellular locations and candidate functions of the SARS-CoV-2 viral proteins studied and provide a resource for future studies of pandemic coronaviruses.

SARS-CoV-2, as the etiological agent of COVID-19, joins SARS-CoV-1 and MERS as an important coronavirus pathogen. Very minor mutations in the viral spike protein (55–58) along with a number of animal reservoirs in endemic regions represent a significant risk for new pandemic coronavirus strains to emerge (59), underscoring the need to understand coronaviral accessory protein functions and virus-host interactions. Comparative studies that analyze multiple coronaviruses (18, 60), including both highly pathogenic and nonpathogenic, will be very beneficial to understanding what can identify new possibly pandemic virus strains. Such resources may allow the research community to not only address current concerns and also provide insight to address with newly emerging coronaviruses in the future. Currently, the ability of S proteins capable of binding to human ACE2 receptor, such as in SARS-CoV-1, SARS-Cov- 2, and MERS, has been used as an indicator of human pathogenicity. But there are coronaviruses, such as HCoV-NL63, that also use ACE2 as a receptor but only cause mild disease (61). Thus, comparing the actual molecular interactions and effects of viral proteins on the host between pathogenic and non-pathogenic virus strains may provide actual insight on what makes certain coronaviruses more medically dangerous and highlight critical virus-host interactions that may be targeted to reduce disease.

The viral envelope of SARS-CoV-2 must contain the proper structural components comprised of S, E, M, and N with a completed viral genome (62) and transcription of both subgenomic and genomic RNA occurs in membranous compartments (63). Accordingly, coronaviruses devote substantial portions of its large genome to manipulating host processes involved in ER-Golgi transport and endocytic and exocytic activity, which was captured in the proximal interactome. We also found evidence of the interaction of SARS-CoV-2 with MHC class I molecules with M, ORF7a, ORF7b, ORF8, and ORF10. Down-regulation of surface expressed proteins has been reported for SARS (64). It is still an open question to how SARS-CoV-2 affects surface expression of important host receptors, which viral proteins affect this process, and the effects on virus replication and disease.

Translation inhibition is a general strategy utilized by many virus families including other RNA viruses like orthomyxoviruses (65), picornaviruses (66, 67), rhabdoviruses (68), and togaviruses (69). Host translational blockade may broadly block antiviral responses and can also cause affect the viability of the infected cell. Some but not all viruses have strategies to overcome translational shutoff, biasing translation of viral mRNA, including the use of IRES elements (70). Lung tissue from COVID patients, in particular, displayed proteomic changes associated with translation inhibition. NSP1 from both SARS-CoV-1 (9) and SARS-CoV-2 (28, 29) have been shown to be potent inhibitors of host translation and are thought to do so using at least two mechanisms: binding to and inhibition of EIF3 translation initiation complex and direct cleavage of host mRNAs. Cryo-EM studies place a domain of NSP1 as sitting in the mRNA channel of the 40S ribosome. Our proximity proteomics data shows NSP1 of SARS-CoV-2 binding to a significant number of EIF3 complex subunits and we demonstrate that NSP1 is able to block translation of capped transcripts as well as transcripts containing host and viral IRES elements. We also observe, as another study has shown (28), that NSP1-induced translational shutoff affects host and viral transcripts containing the viral 5’ UTR. This element exists on all genomic and subgenomic viral RNAs (71). Whether other SARS-CoV-2 factors are necessary to overcome NSP1 translational inhibition or if, during viral replication, the large number of viral transcripts simply outcompetes host transcripts, as seen in vesicular stomatitis virus (72), remains to be determined. A recent study(73) from autopsies of COVID patients characterized whole proteome changes in multiple organs.

Innate immune signaling is a central mechanism of host cell response to viral infection. ORF6 of SARS-CoV-1 (11) and SARS-CoV-2 (38) were shown to be potent inhibitors of such antiviral signaling. One proposed mechanism is that ORF6, through association with specific NPCs (RAE1-Nup98), blocks import of activated transcription factors needed to induce *IFNB1* transcripts and other primary interferon-stimulated genes. In this regard, we identified MAVS proximal to ORF6 and ORF9b. We observed that ORF6, but not ORF9b, inhibited RLR signaling downstream of RIG-I RNA-binding. Taken with the observed adjacencies to nuclear pore proteins noted above, it is likely that the model suggested for ORF6 from SARS-CoV-1 (39) may also be operative for SARS-CoV-2 and that disruption of nuclear import may not be specific only to immune-specific transcription factors but may affect a wider variety of imported proteins.

Viral proteases, such as SARS-CoV-2 NSP5 studied here, have been shown to be potent antiviral targets (74). These proteases are essential for viral replication and escape has proven difficult in resistance studies (75). Coronaviruses encode two proteases NSP3 and NSP5, with NSP5 classified as the main protease. They are both necessary for the processing of the ORF1ab polyprotein containing the viral replicase proteins. NSP5 shows similarity to proteases found in picornaviruses and noroviruses (76). Beyond their importance in viral replication, these viral proteases can target host proteins containing their target residues (77). NSP5 recognizes certain glutamine-serine/alanine/glycine residues, with added specificity being determined by two to three flanking residues (48). Picornavirus virulence has been shown to be mediated in part by 3C protease cleavage of host proteins (44). Using both BioID and SILAC metabolic labeling followed by mass spectrometry, we sought to identify candidate host proteins and use a modified FRET-based cleavage assay to determine if these candidates contained sequences cleavable by NSP5. We identified human CWC22 and FANCD2 as candidates; both proteins contained sequences that could be cleaved by NSP5 in an assay used here which can be used to rapidly assess other potential host targets. The proteomic studies also identified clusters of host factors involved in DNA damage and repair and RNA splicing. Furthermore, we show the effects of GC376 (78), a protease inhibitor of feline coronavirus, displays evidence of inhibition of NSP5 cleavage activity. Consistent with this, GC376 has been shown to block viral replication of SARS-CoV-2 in early studies (79) and we observe that this protease inhibitor blocks NSP5 cleavage of both host and viral target peptide sequences.

The global impacts of the SARS-CoV-2 pandemic have focused attention on identifying new treatments and interventions. Given both the newness of the virus and the relative dearth of research into human coronaviruses, it is important that many resources are generated to better understand aspects of the virus-host interaction. The present work contains a proximal proteomic resource for 17 SARS-CoV-2 viral proteins and combining such proximal proteomics with TAP-based proteomics may be helpful in leveraging the strengths associated with each technique. While proximity proteomics can identify transient, indirect, and weak binding events, including those dependent on intact membranes, TAP-based approaches can focus attention on complexes of proteins that stably associated with each other. We validate the quality of the present proximity data set by corroborating spatial insights with biochemical fractionation experiments. Taken together with other efforts to generate high-quality resources, these data should prove helpful in both generating hypotheses and better understanding dynamics of virus-host interactions in regards to human disease.

## Acknowledgements

This work was supported by the USVA Office of Research and Development, NSF Graduate Research Fellowship 1656518, and by NIAMS/NIH AR45192 (P.A.K.). We thank members of the Wang and Khavari labs for helpful discussions and KMF for her revisions.

## Author Contributions

Conceptualization, J.M.M., M.R., P.A.K.; Methodology J.M.M., M.R., R.L.S.; Formal Analysis, L.D., I.F., M.G.G., X.Y., Y.Z.; Investigation, J.M.M., W.M., M.R., D.S.R., D.R., R.L.S., X.Y., Y.Y.; Writing – Original Draft, J.M.M.; Writing – Review & Editing, J.M.M., M.R., R.L.S., Y.W., P.A.K.; Visualization, L.D., I.F., M.G.G., R.L.S.; Project Administration, J.M.M.; Supervision, Y.W., P.A.K.

## Declaration of Interests

The authors declare no competing interests.

### Materials and Methods

#### Cell Culture

HEK293T were obtained from Takara Bio and were cultured on DMEM 10% FBS, 1% Penicillin/Streptomycin and grown at 37C, 5% CO2. For SILAC experiments,(80) the cells were cultured in a medium containing [^13^C_6_,^15^N_2_]-lysine and [^13^C_6_]-arginine for at least 2 weeks to promote complete incorporation of the stable isotope-labeled amino acids. Cells were tested for mycoplasma prior to experiments using MycoAlert Mycoplasma Detection kit (Lonza).

#### Transfection, Biotin Labeling, and Streptavidin Pulldown

All viral expression constructs were obtained from Addgene (16). HA-BASU was cloned in frame with either an N-terminal or C-terminal linker as indicated. For BioID experiments 5x10^6^ HEK293T were plated and transfected with 5ug of each viral expression plasmid. 24 hours post transfection, biotin was added (50 uM final concentration) for 4 hours, then media was exchanged twice with DPBS and the cells harvested and lysed in RIPA buffer (Thermo Scientific) supplemented with protease inhibitors (. Lysates were sonicated and then, using the Kingfisher Flex automated Purification, incubated for six hours with 100 uL of ReSYN (ReSYN Biosciences) streptavidin microparticles and then washed sequentially with 2% LDS buffer, Triton X-100 buffer (1% Triton X-100 0.1%, Sodium Deoxycholate 500mM, 1mM EDTA, 50mM HEPES pH 7.5), Igepal Wash Buffer (0.5% Igepal, 0.5% Sodium Deoxycholate, 10mM TRIS pH 7.5, 333.3mM LiCL, 20mM EDTA), and deposited into 50mM TRIS pH 7.4. Samples were washed with automated mixing for 30 minutes for each step. A portion of the whole cell lysate was saved and ran on SDS-PAGE gel, transferred to PVDF, and then probed with anti-HA antibody with 800CW anti-rabbit (LICOR) secondary along with 800CW streptavidin dye (LICOR) to confirm viral protein expression and total biotinylation. For SILAC experiments with NSP5 and C145A, approximately 2×10^6^ cells were harvested, washed with ice-cold PBS for three times, and lysed by incubating on ice for 30 min with CelLytic M (Sigma) cell lysis reagent containing 1% protease inhibitor cocktail. The cell lysates were centrifuged at 7,000g and at 4°C for 15 min, and the resulting supernatants collected.

#### Sample Preparation for Mass Spectrometry

After wash and purification samples contained bound proteins on beads in TRIS buffer. The protein on the beads were reduced with dithiothreitol, and alkylated with iodoacetamide. The processed proteins were subsequently digested with Trypsin/Lys-C (Promega) at an enzyme/substrate ratio of 1:100 in 50 mM NH_4_HCO_3_ (pH 8.5) at 37 °C for overnight.

For SILAC samples, the protein lysates prepared from cells with WT or mutant NSP5 were combined at 1:1 ratio (by mass), and 30 µg of the mixed protein lysate was loaded onto a 10% SDS-PAGE gel. After electrophoresis, the gel lanes were cut into 11 slices according to apparent molecular weight ranges of proteins (< 20, 20-25, 25-30, 30-37, 37-42, 42-50, 50-62, 62-75, 75- 100, 100-150, >150 kDa), reduced in-gel with dithiothreitol, and alkylated with iodoacetamide. The processed proteins were subsequently digested in-gel with Trypsin/Lys-C (Promega) at an enzyme/substrate ratio of 1:100 in 50 mM NH_4_HCO_3_ (pH 8.5) at 37 °C for overnight. Subsequently, peptides were recovered from gels with a solution containing 5% acetic acid in H_2_O and then with a solution containing 2.5% acetic acid in an equi-volume mixture of CH_3_CN and H_2_O.

All the resulting peptide mixture was subsequently dried in a Speed-vac, and desalted by employing OMIX C18 pipet tips (Agilent Technologies, Santa Clara, CA). LC-MS/MS experiments were conducted on a Q Exactive Plus mass spectrometer equipped with an UltiMate 3000 UPLC system (Thermo Fisher Scientific).

#### LC-MS/MS Analysis

Samples were automatically loaded at 3 µL/min onto a precolumn (150 µm i.d. and 3.5 cm in length) packed with ReproSil-Pur 120 C18-AQ stationary-phase material (5 µm in particle size, 120 Å in pore size, Dr. Maisch). The precolumn was connected to a 20-cm fused-silica analytical column (PicoTip Emitter, New Objective, 75 µm i.d.) packed with 3 µm C18 beads (ReproSil-Pur 120 C18-AQ, Dr. Maisch). The peptides were then resolved using a 180-min gradient of 2-45% acetonitrile in 0.1% formic acid, and the flow rate was maintained at 300 nL/min.

The mass spectrometer was operated in a data-dependent acquisition mode. Full-scan mass spectra were acquired in the range of *m/z* 350-1500 using the Orbitrap analyzer at a resolution of 70,000 at *m/z* 200. Up to 25 most abundant ions found in MS with a charge state of 2 or above were sequentially isolated and collisionally activated in the HCD cell with a normalized collision energy of 28 to yield MS/MS.

#### Database Search

Maxquant, Version 1.5.2.8, was used to analyze the LC-MS and MS/MS data for protein identification and quantification.(81) The database we used for search was human IPI database, version 3.68, which contained 87,061 protein entries. The maximum number of miss-cleavages for trypsin was two per peptide. Cysteine carbamidomethylation and methionine oxidation were set as fixed and variable modifications, respectively. The tolerances in mass accuracy were 20 ppm for both MS and MS/MS. The maximum false discovery rates (FDRs) were set at 0.01 at both peptide and protein levels, and the minimum required peptide length was 6 amino acids. Spectral match assignment files were collapsed to the gene level and false positive matches and contaminants were removed. SAINT analysis (Choi et al., 2011) (crapome.org) was run with the following parameters: 10,000 iterations, LowMode ON, Normalize ON and the union of MinFold ON and OFF. Minimum interactome inclusion criteria were SAINT≥ 0.9, fold change over matched cell type control ≥ 4. Low normalized spectral count proteins were removed.

#### Gene Ontology

Gene Ontology (GO) term analyses were produced using the clusterProfiler (Yu G. et al., 2012.) R package. Proteins with SAINT score ≥0.9 were classified as likely interactions and used to identify enriched GO terms for the individual SARS-CoV2 protein interactomes. Highly redundant GO terms were removed for readability. Bar plots and heatmaps were produced with the ggplot2 (Wickham H et al. 2016) and heatmap (Kolde R., 2018) packages respectively in R.

#### Host-Virus Interaction Network

Host-virus interaction network produced from BASU BioID interactions with a SAINT score ≥ 0.9 in Cytoscape (Shannon et al. 2003). The network was further curated to emphasize the significantly enriched GO terms for each SARS-CoV-2 protein. Edges denoting Host-virus protein interactions are indicated in red. Host-host interactions were determined from high confidence (>0.700) STRING database interactions obtained from experimental evidence and database interactions for all of the curated proteins. Cell endogenous protein interactions are denoted by grey edges. Clusters were highlighted based on highly enriched GO terms for SARS-CoV-2 proteins.

#### Cellular Fractionation

Cellular fractionations were generated using a previous protocol(82) with minimal modification. Cells were transfected as described previously. Cell pellets were split into three separate samples. The first was lysed using RIPA buffer (Thermo Scientific) and was labeled whole cell lysate (WCL). The second sample was resuspended in buffer containing 0.3% Igepal, 10mM HEPES, 10mM KCl, 1.5mM MgCl_2_. Sample was pelleted at 1500G and supernatant was collected and labeled cytoplasm/membrane fraction. The remaining pellet was washed once and then lysed in RIPA and labeled nuclear fraction. The third sample was lysed in buffer containing 100ug/mL Digitonin, 50mM HEPES, and 150mM NaCl. Sample was pelleted at 2400G and supernatant was collected and labeled cytoplasm fraction. The remaining pellet was washed once and lysed in RIPA and labeled nuclear/membrane fraction. Equal volumes of each fraction along with 20ug of WCL were loaded and ran in a 4-12% Tris-Bis Polyacrylamide Gel (Invitrogen). Samples were transferred to PVDF and blotted for HA (Viral Proteins), Alpha tubulin (cytoplasm control), calnexin (Membrane control), Histone H3 (nuclear control).

#### GWAS COVID Risk SNP Analysis

COVID GWAS datasets were sourced from COVID-19 Host Genome Initiative (https://www.covid19hg.org/), the Ellinghaus, Degenhardt, et al study (21), and from the UK Biobank (https://grasp.nhlbi.nih.gov/Covid19GWASResults.aspx). GWAS hits were converted to hg19 coordinates and phenotypes for each GWAS study were noted. Gene locations are sourced from gencode v19 exon coordinates. The GWAS SNPs were then expanded by LD r2 > 0.8 with phase 1000 Genomes LD information using LDlinkR (83), and phase 1 1000 Genome LD information using HaploReg (84). The expanded SNP list was then overlapped with GTEx lung, spleen, blood, cis-eQTL data, DICE cis-QTL data, and eQTLGen cis-eQTL data (23, 85, 86).

#### HiChIP Data Processing and Virtual 4C Visualization

HiChIP all valid pair matrices for GM12878, Naïve T cells, Th17 cells and Treg were downloaded from GEO (GSE101498, (87)). v4C plots were generated from HiChIP valid pair matrices. The interaction profile of a specific 5-kb bin containing the TRIM4 anchor was then plotted in R. H3K27ac ChIP-seq peaks for GM12878, Naïve T cells, Tregs and T helper cells were downloaded from ENCODE as 1d peak sets. FitHiChIP pipeline was used to call loops with 5kb bin, peak-to-all interaction type, loose background, and FDR < 0.01 (88). The merged significant interaction files from FitHiChIP pipeline along with corresponding ATAC–seq profiles were visualized in WashU Epigenome web browser. Browser shots from WashU track sessions were then included in the v4C and interaction map anecdote.

#### Luciferase Assays

For NSP1 translation assays, in-vitro transcribed transcripts were generated by first PCR amplifying DNA containing T7 promoter followed by UTR or IRES elements and firefly or renilla luciferase. Second, using HiScribe™ T7 ARCA mRNA Kit (with tailing) (NEB) capped and polyadenylated transcripts were synthesized. 5x10^5^ 293T cells were transfected with 2ug of plasmids expressing either GFP or NSP1 and then incubated overnight. The next day the cells were transfected with 2ug of the corresponding IVT transcripts and were harvested 8 hours post second transfection. Cells were harvested with 400ul of Passive Lysis Buffer (Promega) and quadruplicate samples were plated on an opaque 96 well plate. 50ul of LARII firefly luciferase substrate (Promega) was added and the plate was read on the luminescence setting of the Spectramax i5 plate reader. 50ul of Stop & Glo renilla luciferase substrate was then added and the plates reread. For *IFNB1* promoter activity assays, 2.5x10^5^ 293T cells were transfected with 2ug of plasmids expressing either ORF6 or ORF9b along with 1ug of plasmid containing nanoluciferase under the control of a the human *ifnb1* promoter and 50ng of a plasmid containing firefly luciferase under the control of the constitutive TK promoter and either 1ug of empty vector or 1ug of a plasmid expressing the 2-CARD domain of RIG-I. 24 hours post transfection cells were harvested in 200ul of Passive Lysis Buffer and triplicate samples were plated on an opaque 96 well plate. Nano-Glo Dual-Luciferase Reporter Assay System (Promega) was used to obtain a firefly luciferase reading for IFN-beta promoter activity normalized to the firefly luciferase transfection control.

#### NSP5 Cleavage Site Prediction

Protein sequences for hits from SILAC and BASU-BioID proteomics experiments were run through the NetCorona algorithm (48) using the web application: (https://services.healthtech.dtu.dk/service.php?NetCorona-1.0). For Coronavirus Polyprotein controls, the SARS-COV-2 ORF1ab protein sequence (from Uniprot Fasta UP000464024) was run through the NetCorona web application. A previously tested SARS-COV-1 sequence(48), VATLQAENV, was found to be shared in the SARS-COV-2 protein sequence and was also used as a control.

#### FRET-based NSP5 Cleavage Assay

Predicted NSP5 cleavage sites were cloned into ECFP-TevS-YPET (Addgene Plasmid #100097)(89) Briefly, the plasmid was re-cloned to put the Tev Protease Site between an XbaI site and a BsiWI site. Cleavage sequences were cloned in between XbaI and BsiWI sites, restoring the XbaI and BsiWI sites, with one Glycine on each side of the predicted NSP5 cleavage sequences. This approach was based on a cloning strategy used previously to study norovirus protease cleavage sites(90). For protease cleavage assay, 3e4 HEK 293T cells were plated in DMEM + 10% FBS into 96 well black, clear bottom plates (Greiner). 24 hours later, cells were transfected in quadruplicate with 0.1ug of FRET plasmid containing the NSP5 cut-site and 0.1ug of either WT-NSP5 or mutant NSP5^C145A^ expression plasmids (16) with Lipofectamine 3000 following manufacturer protocol for 96-well plates. 24 hours later, media was removed and PBS was added to wells, and wells were imaged on a Spectramax i5 instrument (Molecular Devices) with the following wavelengths: 420/485 nm for ECFP, 485/535 nm for YPET and 420/535 nm for FRET as previously described(89). After background subtraction of un-transfected wells, FRET efficiency was calculated as FRET/ECFP.

#### Quantification and Statistical Analysis

Gene ontology adjusted p-values were produced using the Benjamini-Hochberg method. For heatmaps, a threshold was set whereby at least one protein had a significant score for the presented GO terms. A -Log10 p-Value threshold of >5 for M proteins and >1.3 for all others was used for ‘non-stringent’ heatmaps (with the exception of molecular function heatmaps, which uses an M protein threshold of 3) and -Log10 p-Value > 5 for M proteins and >3 for all others was used for more stringent heat maps presented in figures 1D and 2A

Graphed data are expressed as mean ± SEM and sample size (N) represents independent experiments as noted. Statistical analysis was performed in GraphPad Prism 7 and described in the Fig. legends.

Student’s t test was performed comparing the means between GFP controls and the experimental conditions where N is three independent experiments. For IFNb1 reporter assays, empty vector without RIG-I 2-CARD was set to one and all other conditions are relative to that empty control and is the average of three independent experiments.

NSP5 FRET-cleavage was calculated as described previously and Student’s t test was performed comparing the means between NSP5^C145A^ mutant and NSP5 WT where N is three independent experiments.

Plasmids used in this study

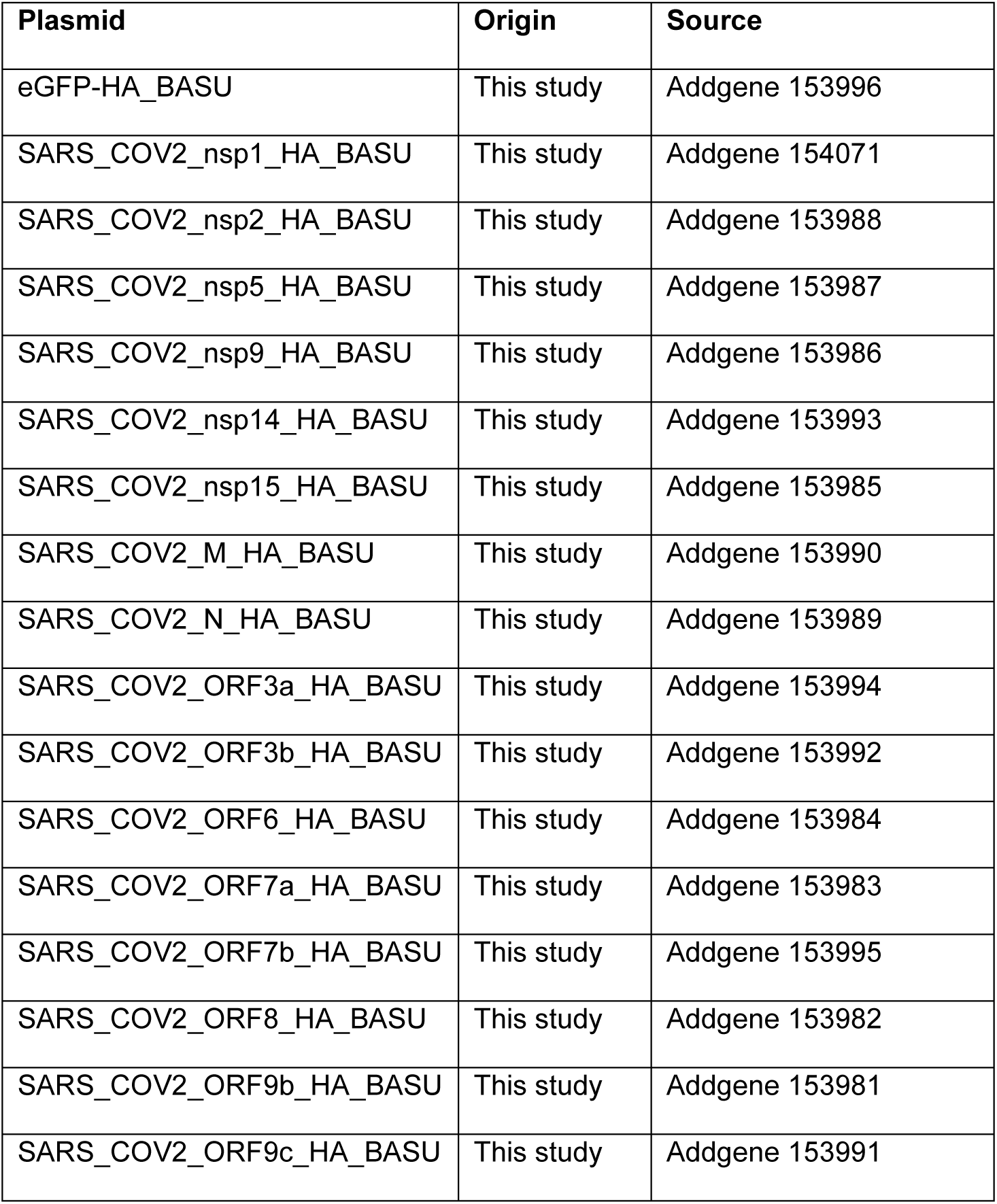
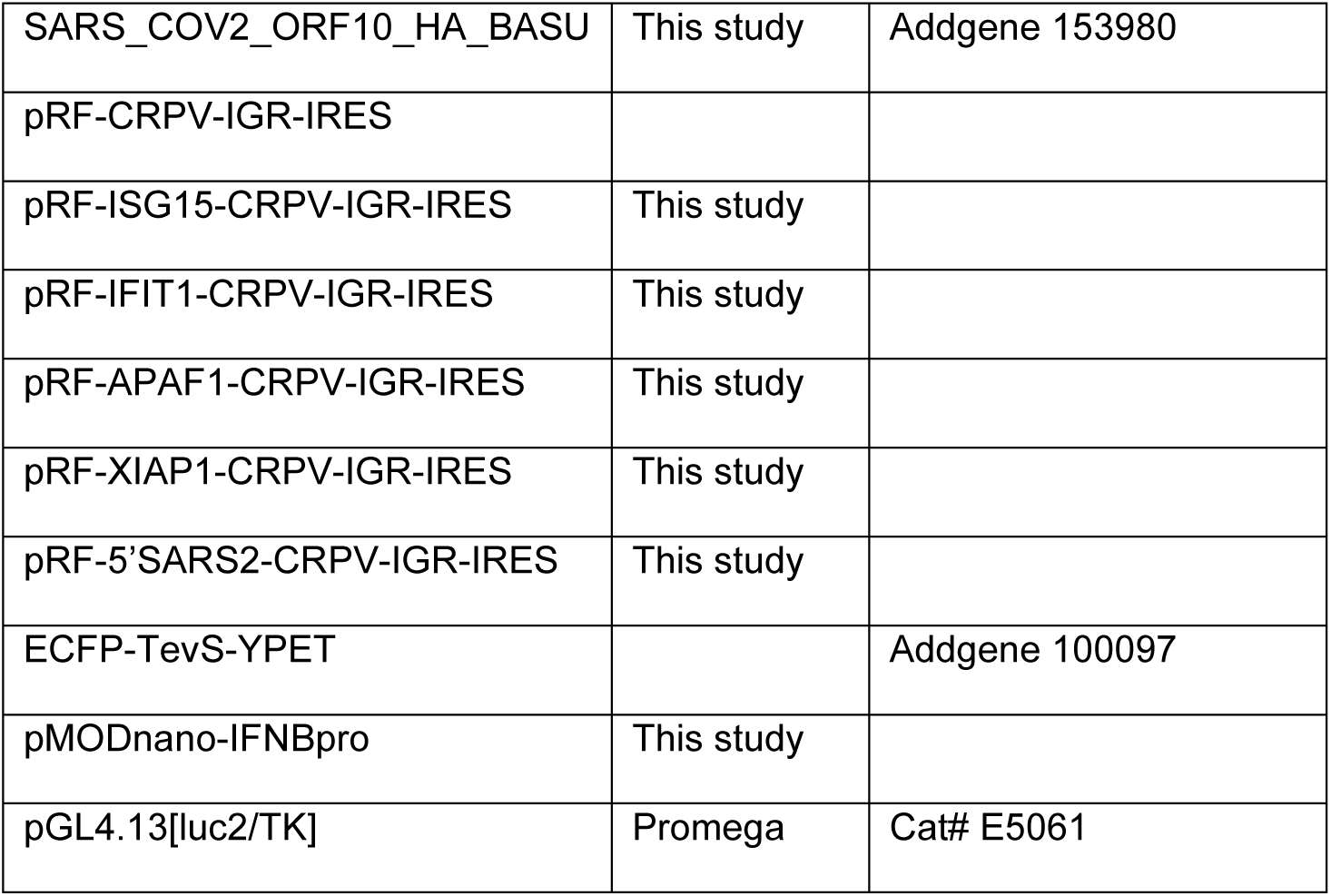

| Plasmid | Origin | Source |
| --- | --- | --- |
| eGFP-HA_BASU | This study | Addgene 153996 |
| SARS_COV2_nsp1_HA_BASU | This study | Addgene 154071 |
| SARS_COV2_nsp2_HA_BASU | This study | Addgene 153988 |
| SARS_COV2_nsp5_HA_BASU | This study | Addgene 153987 |
| SARS_COV2_nsp9_HA_BASU | This study | Addgene 153986 |
| SARS_COV2_nsp14_HA_BASU | This study | Addgene 153993 |
| SARS_COV2_nsp15_HA_BASU | This study | Addgene 153985 |
| SARS_COV2_M_HA_BASU | This study | Addgene 153990 |
| SARS_COV2_N_HA_BASU | This study | Addgene 153989 |
| SARS_COV2_ORF3a_HA_BASU | This study | Addgene 153994 |
| SARS_COV2_ORF3b_HA_BASU | This study | Addgene 153992 |
| SARS_COV2_ORF6_HA_BASU | This study | Addgene 153984 |
| SARS_COV2_ORF7a_HA_BASU | This study | Addgene 153983 |
| SARS_COV2_ORF7b_HA_BASU | This study | Addgene 153995 |
| SARS_COV2_ORF8_HA_BASU | This study | Addgene 153982 |
| SARS_COV2_ORF9b_HA_BASU | This study | Addgene 153981 |
| SARS_COV2_ORF9c_HA_BASU | This study | Addgene 153991 |
| SARS_COV2_ORF10_HA_BASU | This study | Addgene 153980 |
| pRF-CRPV-IGR-IRES |  |  |
| pRF-ISG15-CRPV-IGR-IRES | This study |  |
| pRF-IFIT1-CRPV-IGR-IRES | This study |  |
| pRF-APAF1-CRPV-IGR-IRES | This study |  |
| pRF-XIAP1-CRPV-IGR-IRES | This study |  |
| pRF-5'SARS2-CRPV-IGR-IRES | This study |  |
| ECFP-TevS-YPET |  | Addgene 100097 |
| pMODnano-IFNBpro | This study |  |
| pGL4.13[luc2/TK] | Promega | Cat# E5061 |

Antibodies Used in This Study

| Antibody | Origin | Catalog Number |
| --- | --- | --- |
| HA-tag (C29F4) (Rabbit) | Cell Signaling Technologies | Cat# 3724; RRID:AB_1549585 |
| HA-tag (Mouse) | Abcam | Cat# ab130275; RRID:AB_11156884 |
| HA-tag (Mouse) | Cell Signaling Technologies | Cat # 2367 |
| β-Actin | Sigma-Aldrich | Cat# A1978; RRID:AB_476692 |
| Calnexin | Cell Signaling Technologies | Cat# 2679 |
| Lamin A/C | Cell Signaling Technologies | Cat# 4777 |
| 800CW Goat anti-Mouse IgG (H + L) | LI-COR | Cat# 926-32210; RRID:AB_621842 |
| 800CW Goat anti-Rabbit IgG (H + L) | LI-COR | Cat# 926-32211;<br>RRID:AB_621843 |
| 680RD Goat anti-Mouse IgG (H + L) | LI-COR | Cat# 926-68070;<br>RRID:AB_10956588 |
| 800CW Streptavidin | LI-COR | Cat# 926-32230;<br>RRID:AB_2877131 |
| 680RD Streptavidin | LI-COR | Cat# 926-68079;<br>RRID:AB_2877132 |
| 680RD Goat anti-Rabbit IgG (H + L) | LI-COR | Cat# 926-68071; RRID:<br>AB_10956166 |

#### Data Availability

The mass spectrometry proteomics data for SARS-CoV-2 BIOID and NSP5 SILAC experiments have been deposited to the ProteomeXchange Consortium via the PRIDE(91) partner repository with the dataset identifier PXD023239 andPXD023277 respectively. Raw data has been deposited to Mendeley Data and can be accessed through DOI: 10.17632/mj7jnmvx95.1.

